# Taste Receptor Cells in Mice Express Receptors for the Hormone Adiponectin

**DOI:** 10.1101/335927

**Authors:** Sean M. Crosson, Andrew Marques, Peter Dib, Cedrick D. Dotson, Steven D. Munger, Sergei Zolotukhin

## Abstract

The metabolic hormone adiponectin is secreted into the circulation by adipocytes, and mediates key biological functions including insulin sensitivity, adipocyte development, and fatty acid oxidation. Adiponectin is also abundant in saliva, where its functions are poorly understood. Here we report that murine taste receptor cells express adiponectin receptors, and may be a target for salivary adiponectin. Analysis of a transcriptome dataset obtained by RNA-seq analysis of purified circumvallate taste buds, revealed high expression levels for three adiponectin receptor types. Immunohistochemical studies showed that two of these receptors, AdipoR1 and T-cadherin, are localized to subsets of taste receptor cells. Immunofluorescence for T-cadherin was primarily co-localized with the Type 2 taste receptor cell marker phospholipase β2, suggesting that adiponectin signaling could impact sweet, bitter, or umami taste signaling. However, adiponectin null mice showed no differences in taste responsiveness compared to wildtype controls in brief-access taste testing. AAV-mediated overexpression of adiponectin in the salivary glands of adiponectin null mice did result in a small but significant increase in behavioral taste responsiveness to the fat emulsion Intralipid. Together, these results suggest that salivary adiponectin can effect taste receptor cell function, though its impact on taste responsiveness and peripheral taste coding remains unclear.

## Introduction

Recently, numerous peptides that can function as metabolic hormones, or their cognate receptors, have been detected in saliva and/or in taste receptor cells (TRCs) (Zolotukhin 2013). Of the many peptides present in the oral cavity, several appear to modulate taste-evoked behavioral responses (Dotson *et al*. 2013). For example, both glucagon-like peptide-1 (GLP-1) and glucagon signaling impact behavioral taste responsiveness to sweet stimuli (Elson *et al*. 2010; Shin *et al*. 2008; Takai *et al*. 2015), angiotensin-2 impacts salt taste (Shigemura *et al*. 2013), and peptide YY (PYY) signaling is implicated in the modulation of orosensory responses to lipids (La Sala *et al*. 2013). However, the full impact of peptide signaling on taste signaling remains poorly understood.

The anatomical proximity of salivary-expressed peptides with the peripheral gustatory system provides an opportunity for salivary peptides to impact peripheral taste function. Indeed, we previously reported that salivary PYY can modulate behavioral responsiveness to oral lipid stimuli (La Sala *et al*. 2013). Adiponectin is an anti-inflammatory adipokine primarily secreted from adipocytes into the circulation, where it affects many biological functions such as insulin sensitivity and fatty acid oxidation (Awazawa *et al*. 2011; Villarreal-Molina and Antuna-Puente 2012; Yamauchi *et al*. 2002; Yoon *et al*. 2006). In both plasma and saliva, adiponectin is present in multiple oligomeric forms referred to as low, medium, high, and super high molecular weight, the latter found only in saliva (Bobbert *et al*. 2005; Lin *et al*. 2014). The origin of salivary adiponectin is not entirely clear; while it has been shown in humans to be synthesized in salivary gland ductile cells (Katsiougiannis *et al*. 2006), it is likely that adiponectin is also transferred to saliva from the circulation (Wang *et al*. 2013). The function of salivary adiponectin is debated, though a few studies suggest that it influences saliva secretion and plays an anti-inflammatory role in the oral cavity (Ding *et al*. 2013; Katsiougiannis *et al*. 2010). To our knowledge, no role in gustation has been shown for adiponectin.

By combining RNA-seq analysis of a murine circumvallate (CV) taste bud transcriptome with immunohistochemistry (IHC) in this tissue, we identified two canonical adiponectin receptors – T-cadherin and Adipor1 – expressed in mouse TRCs (Crosson SM, *et al*. submitted for publication). The localization of these receptors to functional subsets of taste cells, along with changes in licking to lipid stimuli upon perturbation of oral adiponectin signaling, suggests a role for salivary adiponectin in the modulation of gustatory function.

## Materials and Methods

### Mice

This study was approved by the Institutional Animal Care and Use Committee (IACUC) at the University of Florida. All procedures were done in accordance with the principles of the National Research Council’s guide for the Care and Use of Laboratory Animals. Mice had *ad libitum* access to food and water, except where otherwise noted, and were housed at 22-24°C with a 14/10 hr light/dark cycle. Wildtype (WT) C57BL/6J mice were bread in-house and B6;129-Adipoq^tm1Chan^/J (APN KO, which contains a null allele of the gene encoding adiponectin) mice were purchased from Jackson Labs. In some experiments, APN KO mice each received a total of 1×10^12^ vg of recombinant AAV vector either bilaterally in the submandibular salivary glands or via the tail vein prior to behavioral testing (for details, see (Katano *et al*. 2006)). DNA was isolated from ear punches of all APN KO mice for genotyping to confirm exon2 deletion in the *Adipoq* gene. Genotyping primers are reported in Table S1.

### Tissue Collection

Mice were deeply anesthetized by i.p. injection of a ketamine/xylazine mixture (200mg/kg and 10mg/kg respectively), then perfused intracardially with 4% paraformaldehyde in phosphate buffered saline (PBS, pH~7.4), followed by tissue dissection. Tissues were fixed overnight in 4% paraformaldehyde in PBS (pH~7.4), cryoprotected by incubation with 30% sucrose in PBS (pH~7.4) overnight, and frozen in O.C.T. mounting medium prior to cryosectioning. Mice used for 5-HT tissue staining were injected with 5-HTP in Lactated Ringers Solution (200 mg/kg) one hour before euthanasia to increase the amount of 5-HT in Type 3 TRCs.

### Immunohistochemistry

#### Adiponectin receptor immunofluorescence

Specific information regarding antibody sources, dilutions and species is located in Table 1. OCT-imbedded tongues were sectioned in 10-20 μm coronal slices using a cryostat (Leica CM3050 S; Leica Microsystems, Nussloch GmbH, Germany) and mounted on Fisher Superfrost Plus slides. Immunohistochemistry was conducted using traditional indirect immunofluorescence. All washing steps were done using TBST (50 mM Tris-HCl, 0.9% NaCl, and 0.5% Tween 20, pH~7.6). Tissues were blocked for one hour at room temperature with in-house blocking buffer (5% normal donkey serum in TBST) to reduce non-specific antibody binding. Sections were then incubated overnight at 4°C with primary antibody diluted in blocking buffer, followed by secondary antibody incubation with either a Donkey-anti-Rabbit IgG Alexa488 conjugate or a Donkey-anti-Goat IgG Alexa649 conjugate (1: 1000 dilution in blocking buffer, one hour at room temperature). All sections were counterstained with 4’,6-diaminidino-2-phenylindole (DAPI) and visualized via a confocal microscopy (Leica SP5).

**Table 1.** Host species, dilution, and supplier information for primary antibodies used in IHC experiments.

| 1° Antibody | Immunogen | Host | Supplier | Dilution |
| --- | --- | --- | --- | --- |
| AdipoR1 | Synthetic peptide sequence corresponding to amino acids 14-32 of human AdipoR1<br>GAPASNREADTVELAELGP | Rabbit | Dr. Xiao-Rong Peng<br>(Bjursell *et al.* 2007) | 1:200 |
| AdipoR2 | Synthetic peptide sequence corresponding to amino acids 11-29 of human AdipoR2<br>CSRTPEPDIRLRKGHQLDG | Rabbit | Dr. Xiao-Rong Peng<br>(Bjursell *et al.* 2007) | 1:200 |
| T-Cadherin | Recombinant human Cadherin-13 | Goat | R&D systems<br>(Minneapolis, MA, U.S.A. AF3264) | 1:500 |
| NTPDase2 | Mouse NTPDase2 | Rabbit | J. Sévigny, Université Laval, Quebec, Canada. #mN2-36I6 | 1:500 |
| PLCβ2 | Synthetic peptide corresponding to the C-terminal region of human PLCβ2 | Rabbit | Santa Cruz Biotechnologies<br>(Dallas, TX, U.S.A. cat No. sc-206) | 1:500 |
| Gα<br>Gustducin | Synthetic peptide corresponding to an internal region of rat Gα Gustducin | Rabbit | Santa Cruz Biotechnologies<br>(Dallas, TX, U.S.A. cat No. sc-395) | 1:500 |
| NCAM | Purified chicken NCAM | Rabbit | Millipore (Temecula, CA, USA; cat. No. AB5032) | 1:500 |
| 5-HT | Serotonin conjugated to BSA | Rat | Millipore (Temecula, CA, USA; cat. No. MAB352 clone YC5/45) | 1:500 |
| Krt8 | Mouse Cytokeratin 8/18 | Rat | University of Iowa Developmental Studies Hybridoma Bank (antibody Registry ID AB_531826) | 1:500 |

#### Double Labeling Immunofluorescence

Double-labeling techniques were used to co-localize T-cadherin with established TRC markers to characterize expression in specific TRC subpopulations. TRC marker information is located in Table 1 along with other primary antibody information. Double labeling experiments used primary antibodies from different host species, and thus utilized a standard indirect dual immunofluorescence staining protocol. Specifically, tissues were incubated simultaneously with both primary antibodies at 4°C overnight, followed by simultaneous incubation with two secondary antibodies for 1 hour at room temperature. Slides were washed with TBST between each incubation to remove excess antibody. Donkey-anti-Rabbit Alexa488 (1:1000), Donkey-anti-Rat Cy3 (1:200) and Donkey-anti-Goat Alexa649 (1:1000) were used as secondary antibodies to detect antisera from each of the three host species used.

### Plasmid Construction

Three plasmid transgene constructs were used to generate the different adiponectin rescue mouse models reported here. The pTR-Acrp30 plasmid, used to produce the AAV8-APN vector, was assembled by ligating mouse adiponectin cDNA into the pTR-UF backbone (Zolotukhin *et al*. 1996), as described in detail by (Shklyaev *et al*. 2003). To generate pTR-GFP-miR, we ligated miR122 and miR206 target site triplicate oligonucleotides, synthesized commercially, into the 3’ untranslated region (UTR) of an inverted terminal repeat (ITR)-containing plasmid backbone using standard cloning techniques. Lastly, to create pTR-APN-miR, the GFP transgene of pTR-GFP-miR was swapped with mouse adiponectin cDNA, amplified from pTR-Acrp30. Expression of each transgene cassette is driven by the ubiquitous chicken β-actin promotor, and all plasmid constructs were confirmed by Sanger sequencing prior to the production of AAV vectors. Primers used for cloning are reported in Table S1.

### AAV Vector Production and Administration

Recombinant AAV vectors were produced in HEK 293 cells using a triple (AAV5) or double (AAV8) transfection method, and purified via Iodixanol density centrifugation as previously described (Zolotukhin *et al*. 1999). For both AAV5 preps, pHelper (Agilent cat no. 240071-52) was used to supply the adenoviral helper genes and pACG2R5C (Zolotukhin *et al*. 2002) was used to supply the AAV2 *rep* and AAV5 *cap* genes. For AAV8 production, a single helper plasmid (pDG8) containing both the adenoviral genes as well as the AAV2 *rep* and AAV8 *cap* genes was used (Grimm *et al*. 1998). Table 2 shows the plasmids used in the transfection to produce each recombinant AAV vector. Vectors were titered using a PicoGreen-based assay described by (Piedra *et al*. 2015), and were sterile filtered before administration to animals.

**Table 2.**
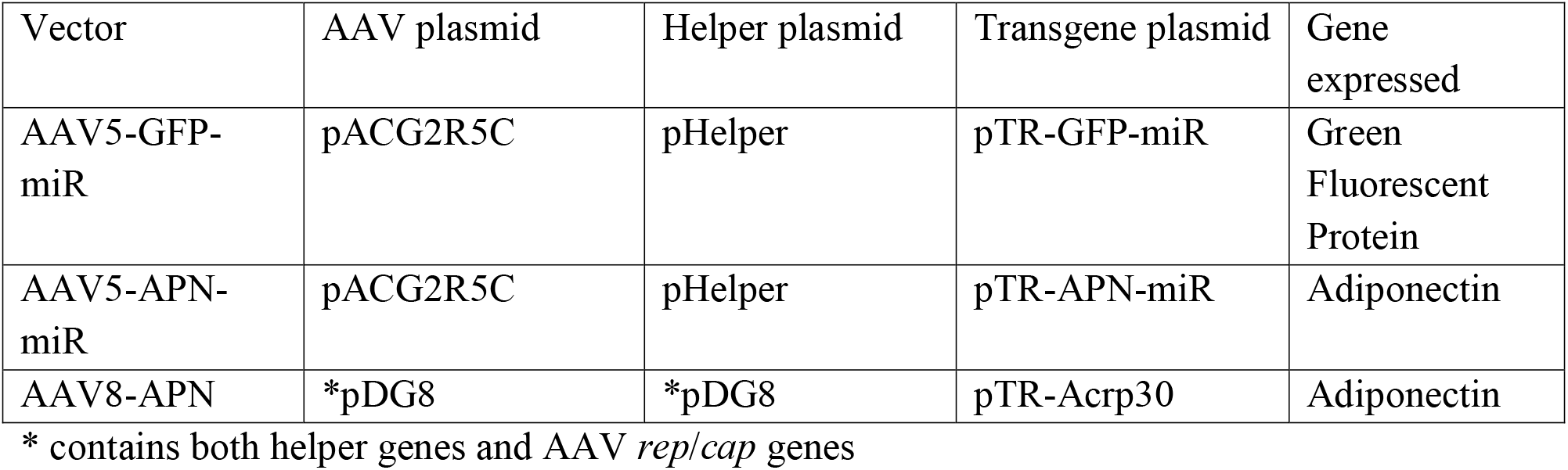
Plasmid constructs used in the production of rAAV vectors.

### Behavior

#### Animals

For behavioral taste testing of APN KO mice and WT mixed background controls (B6129SF2/J), adult mice (10-12 weeks old) were ordered from Jackson Labs, and allowed to acclimate to their new housing environment for 2 weeks prior to behavioral testing. During this two week acclimation period, mice were given *ad libitum* access to food and water up till the start of training/testing, and housed individually. In the second set of behavioral experiments, APN KO mice (10-12 weeks old) were administered either AAV8-APN, AAV5-APN-miR, or AAV5-GFP-miR one month before the first day of training. AAV was administered either via either tail vein injection (AAV8-APN) or submandibular salivary gland cannulation (AAV5-APN-miR and AAV5-GFP-miR) as previously described (Katano *et al*. 2006). After vector administration, mice were single housed and given *ad libitum* access to food and water until the start of the training/testing sessions.

#### Taste Stimuli

All tastants were prepared in 18.2 MΩ ultrapure water and dilutions were prepared fresh before each testing session. Tastants and dilution factors used are listed as follows: citric acid (CA; 0.3, 1, 3, 10, 30, and 100 mM; Sigma-Aldrich), NaCl (30, 100, 200, 300, 600, and 1000 mM; Sigma-Aldrich), quinine hydrochloride (QHCl; 0.03, 0.1, 0.3, and 1, 3 mM; Sigma-Aldrich), sucrose (25, 50, 100, 200, and 400 mM; Fisher Scientific), Intralipid (1.25, 2.5, 5, 10, and 20%; Sigma-Aldrich). Each solution was presented at room temperature and water was used as a “no stimulus” control for each tastant.

#### Procedure

Training and testing procedures were done in a Davis Rig lickometer (Davis MS-160; DiLog Instruments, Tallahassee, FL, USA). The lickometer allows mice access to a sipper bottle containing the stimulus, and uses AC current to record each lick. The lickometer utilizes a motorized table and shutter to restrict the mice to 5 sec trials for each sipper tube. The total session times were 25 min, during which mice could initiate as many trials as they wanted. Mice were tested according to previously published protocols (Elson *et al*. 2010; Glendinning *et al*. 2002; La Sala *et al*. 2013). Two protocols were used; one for appetitive stimuli (sucrose and Intralipid), and one for aversive stimuli (NaCl, CA, and QHCl). For the appetitive stimuli, mice were food and water restricted (1 g food and 2 ml water) for the 23.5 hour period prior to testing. After each testing period, mice were given a 24 hour recovery period where they had *ad libitum* access to food and water. For aversive stimuli, mice were put on a 23.5 hour water restriction schedule throughout the training/testing period and given *ad libitum* access to food. During aversive stimuli testing, a water rinse was presented in between each stimulus presentation in order to control for potential carryover effects. All mice were weighed daily and given 24 hr supplementary *ad libitum* access to food and water if at any time their weight dropped below 85% of their pre testing weight.

#### Data Analysis and Statistics

For aversive stimuli, tastant/water lick ratios were obtained by dividing the average number of licks per trial for each stimulus concentration, by the average number of licks per trial to water. For appetitive stimuli, a standardized lick ratio was used to control for the impact of small changes in water licks. The standardized lick ratio is calculated by dividing the average number of licks per trial for each stimulus concentration, by the maximum potential lick rate for that animal as determined by the mean interlick interval distribution during water spout training (Glendinning *et al*. 2002). This controls for individual differences in maximal lick rates. All ratio scores were analyzed pairwise between groups by two-way analysis of variance (ANOVA). If a significant interaction was observed, a *post hoc* Holm-Sidak t-test was used (p < 0.05) to determine if behavioral responses were significantly different between groups, for each individual concentration. Only mice that initiated at least one trial for every concentration were used in the analysis of a given stimulus. For presentation of behavioral data, curves were fit to the mean data for each group using a 2- or 3-parameter logistic function as previously described (Elson *et al*. 2010).

## Results

Numerous reports have shown that taste responsiveness can be modulated by peptide signaling in taste buds. In a previous study from our group, we found that receptors for the peptide PYY are expressed in subsets of TRCs (Hurtado *et al*. 2012; La Sala *et al*. 2013). We then asked if taste buds express receptors for other peptides enriched in saliva. To initially address this question, we queried the transcriptome recently generated by us using RNA-seq of purified CV taste buds obtained from C57BL6/J mice (Crosson SM, *et al*. submitted for publication).

Transcripts encoding three adiponectin receptors – *Adipor1, Cdh13*, and *Adipor2* – were highly expressed in these taste buds (Figure 1). Average *Adipor1* expression was the highest (138.83 ± 8.89 fpkm), followed by *Cdh13* (93.93 ± 12.69 fpkm), and *Adipor2* (30.62 ± 5.84 fpkm). Several other peptide receptors that have been previously reported in taste buds were also found in this dataset, albeit at lower expression levels than those seen for the adiponectin receptors; these include the insulin receptor *Insr* (Baquero and Gilbertson 2011), oxytocin receptor *Oxtr* (Sinclair *et al*. 2010), GLP-1 receptor *Glpr1* (Martin *et al*. 2009; Shin *et al*. 2008), and neuropeptide Y receptor *Npyr1* (Hurtado *et al*. 2012; Zhao *et al*. 2005).

**Figure 1.**
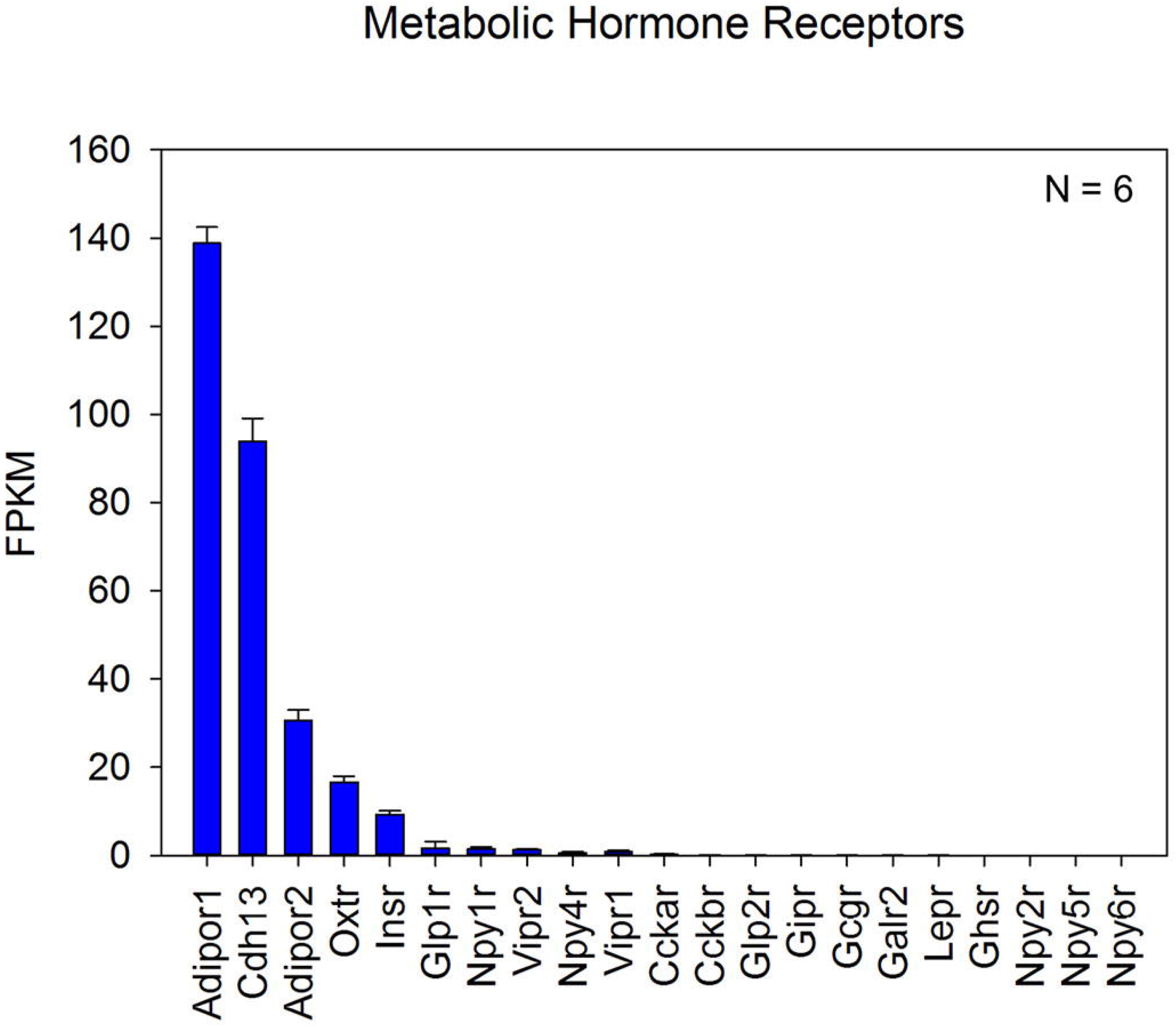
Gene expression levels of metabolic hormone and peptide receptors in WT TRCs. Expression levels for select receptors as determined by RNA-seq of WT murine CV taste buds. The three highest expressing transcripts – *Adipor1, Cdh13*, and *Adipor2* – are all receptors for adiponectin. A total of 6 biological replicates (N = 6) were used for analysis.

To validate the expression of *Adipor1, Adipor2*, and *Cdh13* in mouse TRCs, we performed immunohistochemisty (IHC) on cryosections containing CV papillae from C57BL6/J mice, using polyclonal antibodies against Adipor1, Adipor2, and T-cadherin (encoded by *Cdh13*). Each adiponectin receptor antisera had been previously validated in knockout mouse models (Bjursell *et al*. 2007; Denzel *et al*. 2010). Both Adipor1 and T-cadherin immunolocalize to taste buds (Figure 2A, C). Adipor2 however, does not immunolocalize to taste buds; rather, it is found only in the surrounding tissues (Figure 2B). Co-labeling sections for Adipor2 and cytokeratin 8 (Krt8), a general TRC marker (Mbiene and Roberts 2003), confirmed that taste buds do not express Adipor2 and suggests that the presence of Adipor2 in the RNA-seq database was due to contamination of the taste bud samples with surrounding non-taste tissue.

**Figure 2.**
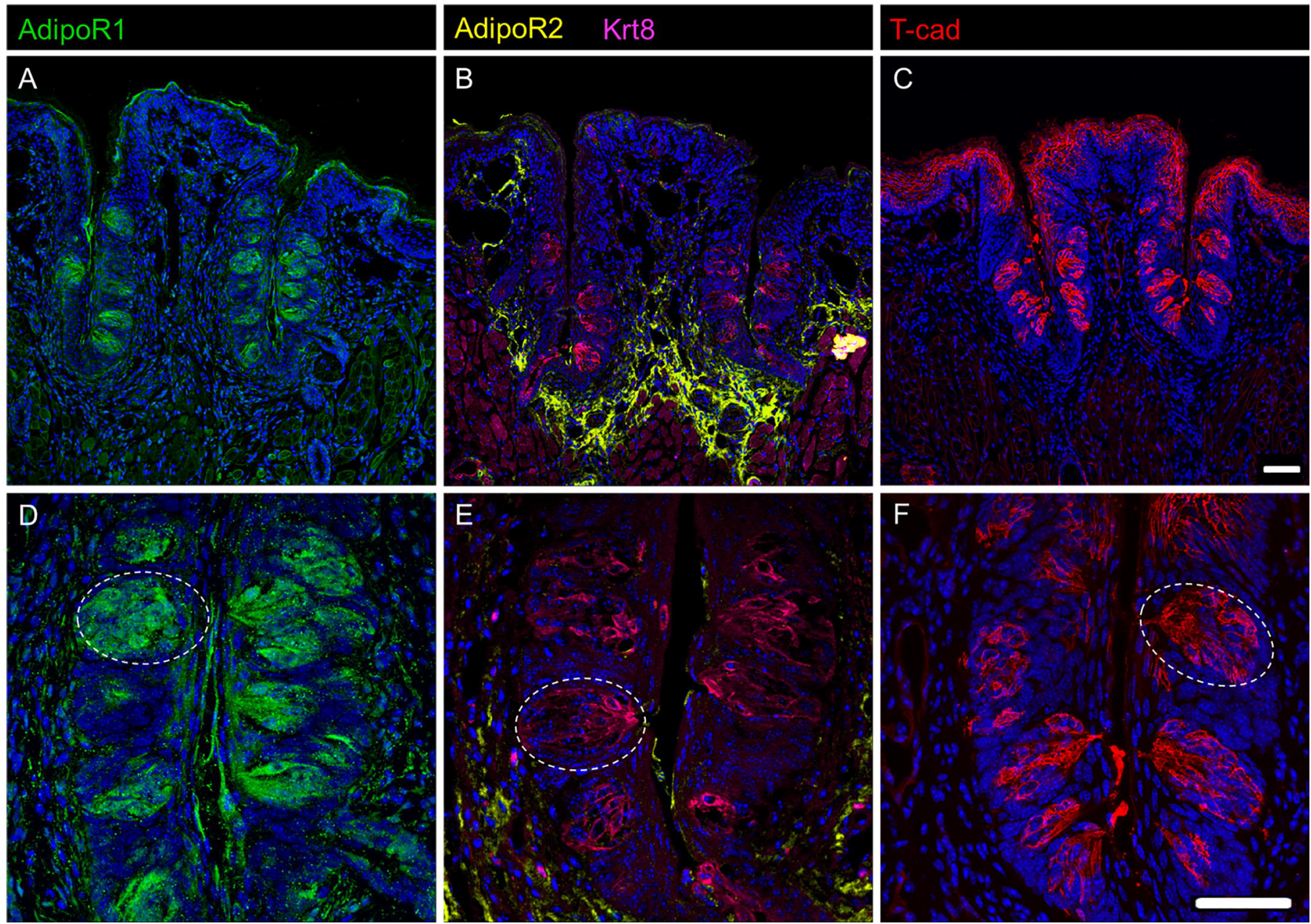
Expression of Adipor1 and T-cadherin in CV TRCs. IHC staining of WT murine CV sections for all three adiponectin receptors, Adipor1 (A, D), Adipor2 (B, E), and T-cadherin (C, F). Adipor1 (A, D) and T-cadherin (C, F) are expressed in CV taste buds (white dotted circle) while Adipor2 (B, E) is expressed in surrounding tissue. Adipor2 sections were costained with Krt8 (B, E), a general taste cell marker. Scale bar is 20 microns.

Mammalian taste buds are composed of multiple TRC types, each of which play different roles in the detection and transmission of taste information (Chaudhari and Roper 2010). To gain insight into the roles of adiponectin signaling in TRCs, we co-localized T-cadherin with established markers for the three main TRC subtypes (Figures 3 & 4): NTPDase2 (Type 1 TRCs, which are thought to play a supporting role; (Vandenbeuch *et al*. 2013)), PLCβ2 and the G protein □-subunit gustducin (sweet, bitter, and/or umami-responsive TRCs; (Ming *et al*. 1999; Miyoshi *et al*. 2001), and 5-HT and NCAM (sour-sensitive Type 3 TRCs; (Huang *et al*. 2008; Yee *et al*. 2001). Host species antibody constraints made dual staining difficult for Adipor1. T-cadherin immunostaining largely colocalized with both PLCβ2 and gustducin, suggesting expression of this adiponectin receptor primarily in a major subset of Type 2 TRCs (Figure 3). T-cadherin was not co-expressed with the Type 3 TRC markers 5-HT or NCAM, though a small subset of NTPDase2-expressing Type 1 TRCs showed some T-cadherin staining (Figure 4). We also measured the co-localization of T-cadherin and these TRC markers by correlation analysis (Costes *et al*. 2004). Consistent with the visual analysis of the IHC co-staining, calculated Pearson’s correlation coefficients (Table 3) indicate that T-cadherin is primarily expressed in Type 2 TRCs.

**Figure 3.**
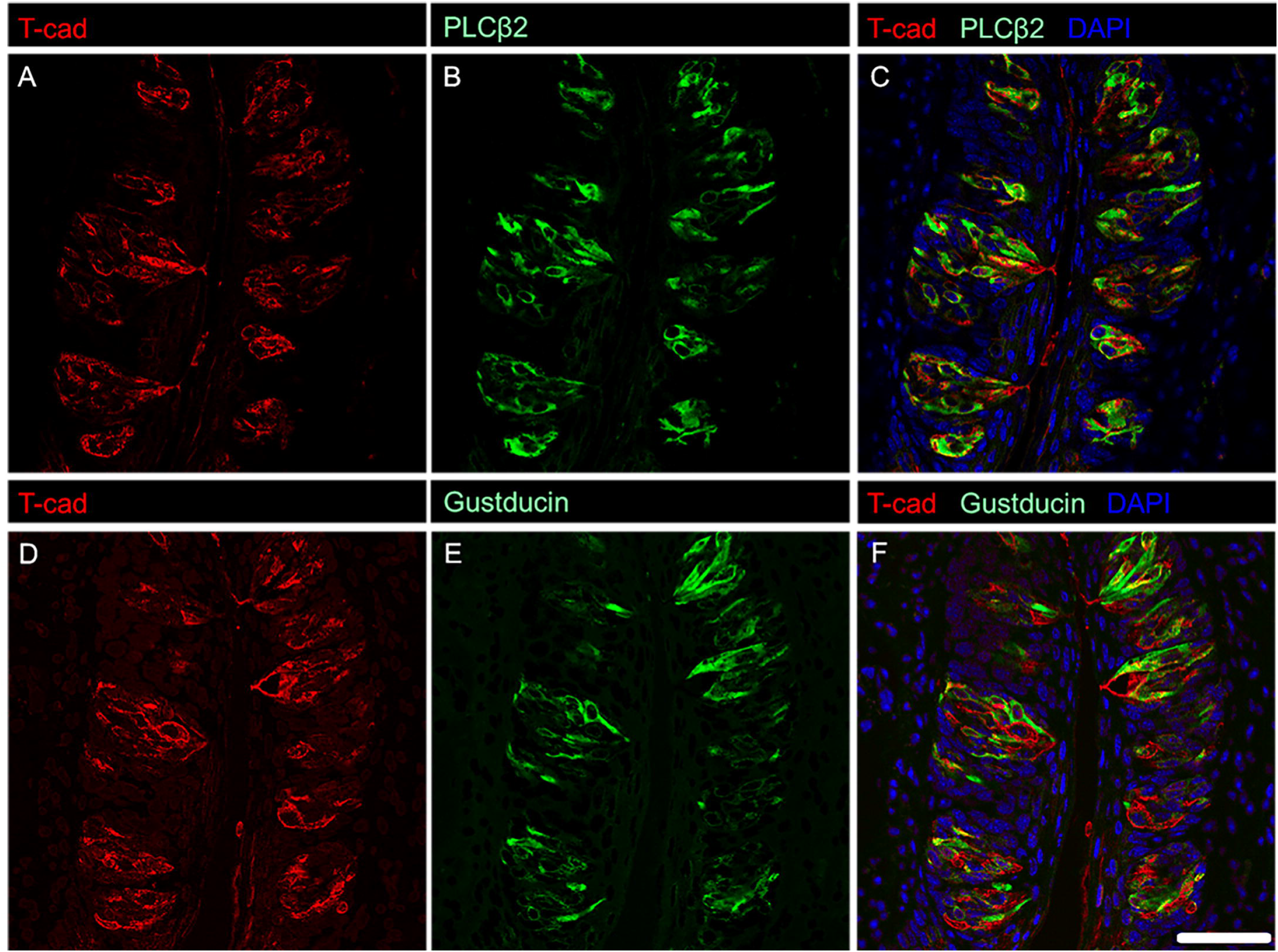
IHC staining of T-cadherin with established markers for sweet, bitter, and/or umami responsive TRCs. T-cadherin localizes to cells expressing PLCβ2 (C), and cells expressing gustducin (F). Single channel images of T-cadherin (A) and PLCβ2 (B) as well as T-cadherin (D) and gustducin (E) are shown for reference. Scale bar is 20 microns.

**Figure 4.**
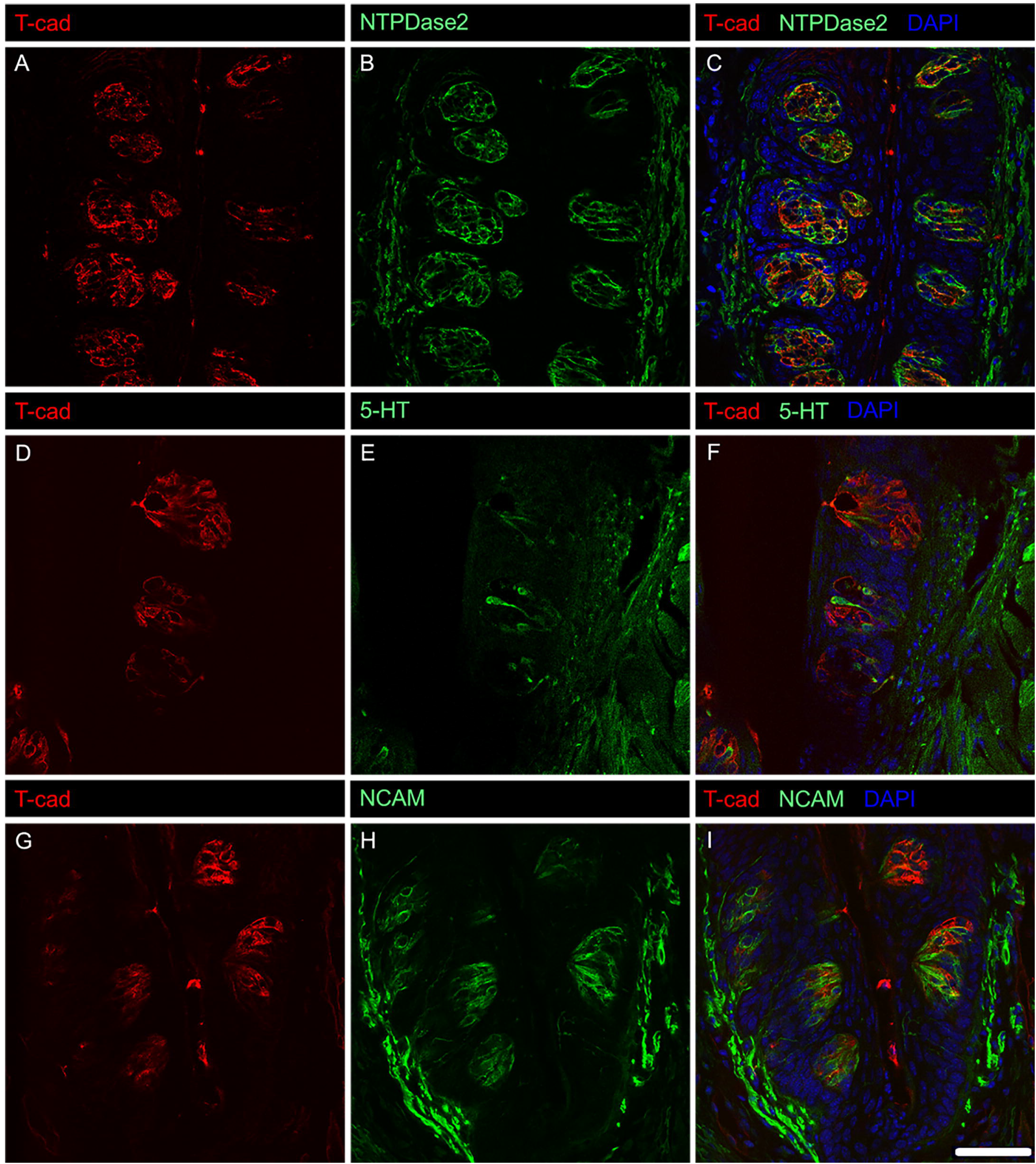
IHC staining of T-cadherin with established markers for supporting and sour responsive TRCs. T-cadherin does not localize to sour responsive TRCs (F, I) and has minimal localization to supporting TRCs (C). Single channel images of T-cadherin (A) and NTPDase2 (B), T-cadherin (D) and 5-HT (E), and T-cadherin (G) and NCAM (H) are shown for reference. Scale bar is 20 microns.

**Table 3.**
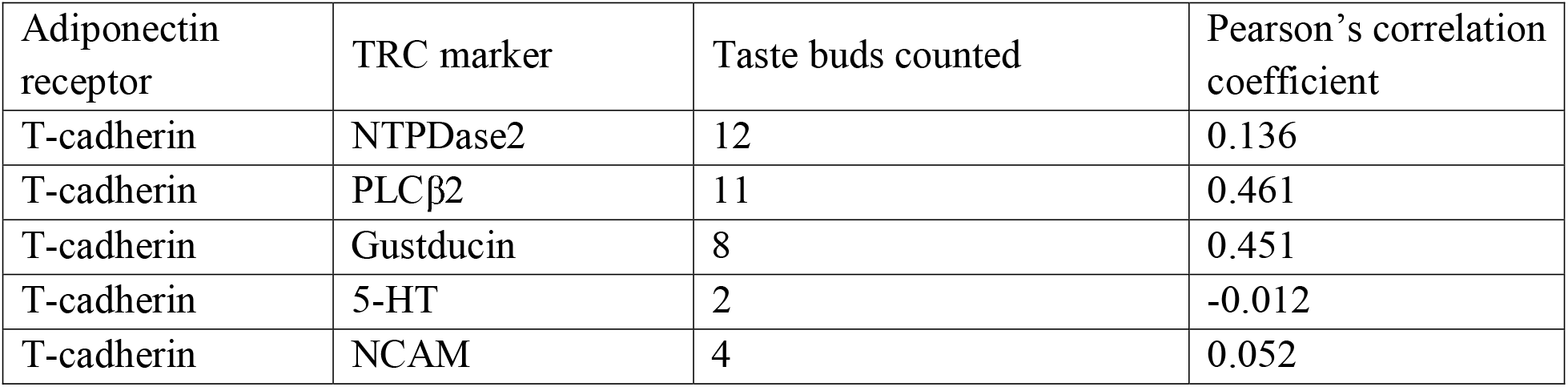
Co-localization analysis of T-cadherin and TRC markers by Pearson’s correlation.

We next asked if adiponectin signaling impacts taste behaviors. We first assessed taste responses in APN KO mice, and their WT controls (B6:129 SF2/J mice) using brief access taste testing. No significant differences in taste responses to sucrose, QHCL, NaCl, CA, or Intralipid were seen between APN KO and control mice (Figure 5).

**Figure 5.**
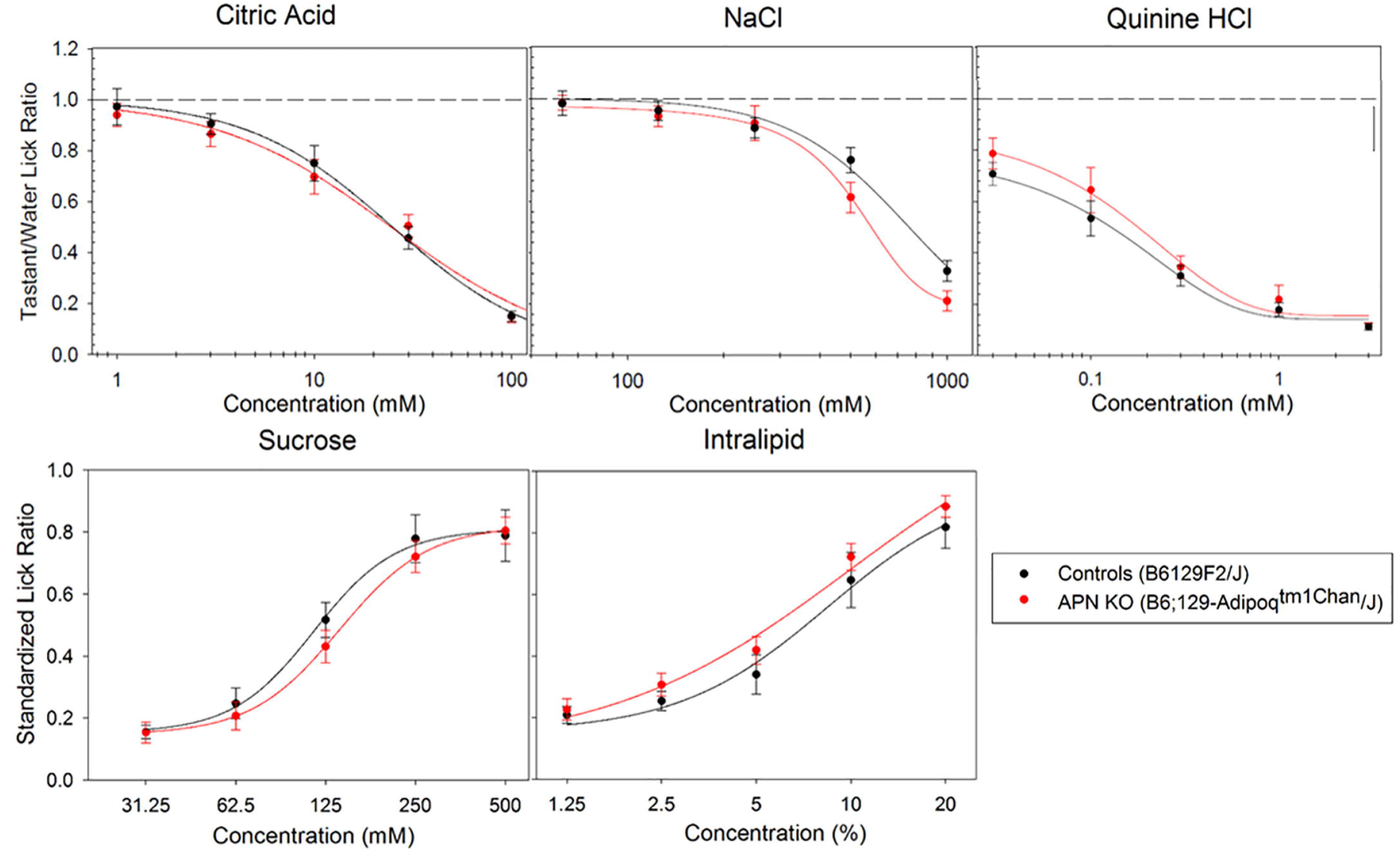
Behavioral taste response comparison of APN KO and WT control mice. Brief-access taste response testing of APN KO (red) and control (black) mice for CA, NaCl, QHCL, sucrose, and Intralipid. No significant difference was observed between groups for any of the stimuli tested, as determined by two-way ANOVA (p > 0.05). A total of 8 mice were used in each group.

To specifically test the effects of salivary and circulating adiponectin, we generated both a salivary gland-specific adiponectin rescue model and, a global overexpressing adiponectin rescue model using recombinant adeno-associated viral (rAAV) vectors in APN KO mice. Since the tissue tropism of AAV serotypes is not well characterized in the salivary gland, we first performed several pilot studies. We chose to focus on AAV serotypes 2, 5, and 8 because AAV2 and AAV5 will reportedly transduce the salivary gland (Katano *et al*. 2006), and AAV8 is a generally robust vector in mice. Mice received a total of 1×10^12^ vg of either AAV 2, 5, or 8 (each expressing a GFP reporter under the chicken β-actin promoter) bilaterally in the submandibular gland (Figure 6). Both AAV5 and AAV8 displayed high salivary gland transduction (Figure 6B,C). However, AAV8 also had high transduction in the liver (Figure 6E), a common off-target tissue for AAVs. Because of the unintended liver transduction observed with AAV8 vectors, we decided to use AAV5 as the vector for our salivary adiponectin rescue and AAV8 as the vector for the global adiponectin rescue. To further increase the specificity of the AAV5 vector for salivary gland transduction, we included micro RNA target sites for miR122 and miR206, which are liver and skeletal muscle specific, respectively (Geisler *et al*. 2013). Using this micro RNA target site containing vector, we were able to abolish off target expression in the liver (Figure 7).

**Figure 6.**
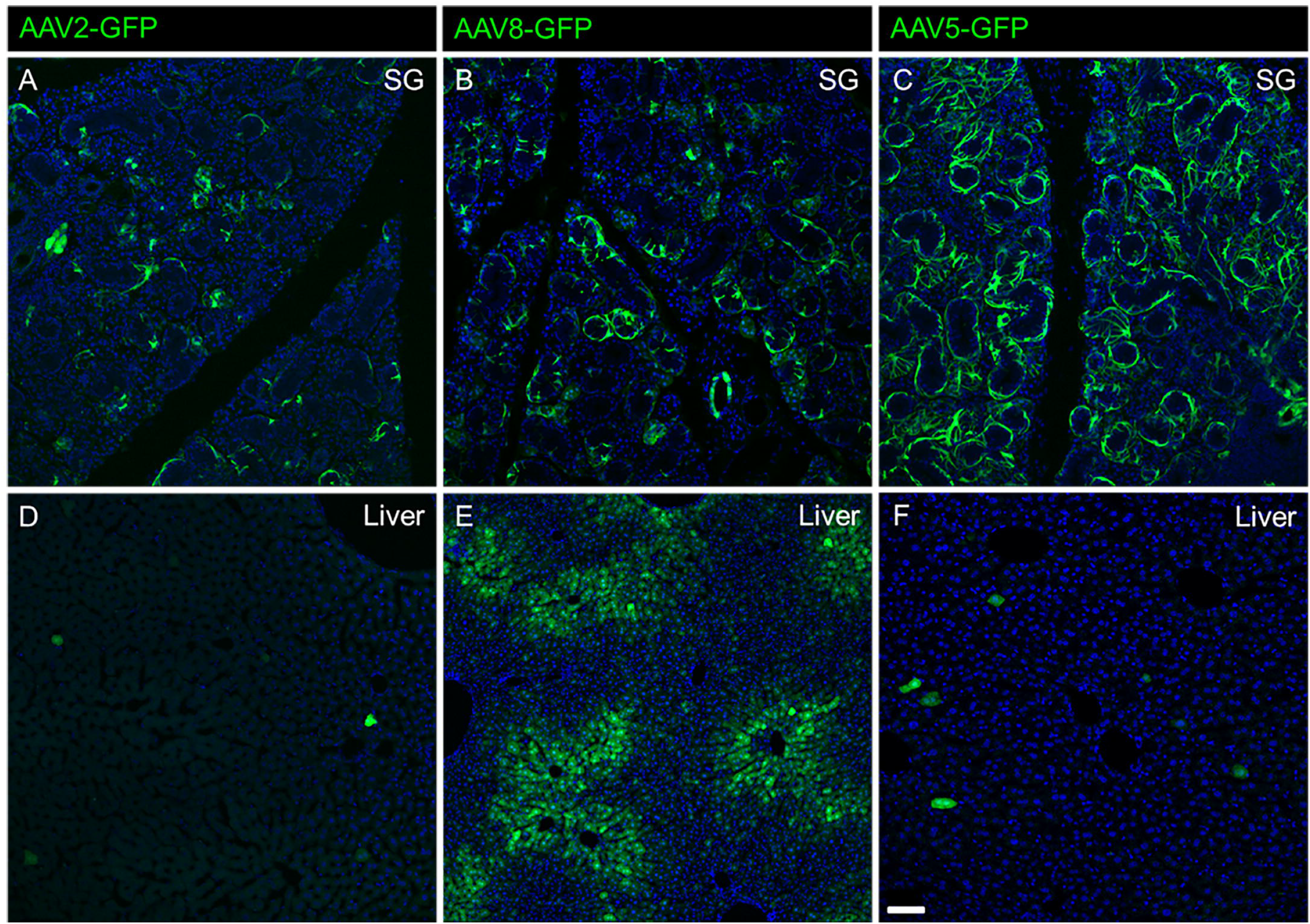
Salivary gland tropism of AAVs 2, 8, and 5. GFP expression in salivary glands (A, B, C) 1 month after AAV2 (A), AAV8 (B), or AAV5 (C) administration in WT mice. Off target liver expression was observed for all vectors AAV2 (D), AAV8 (E), and AAV5 (F). Scale bar is 50 microns.

**Figure 7.**
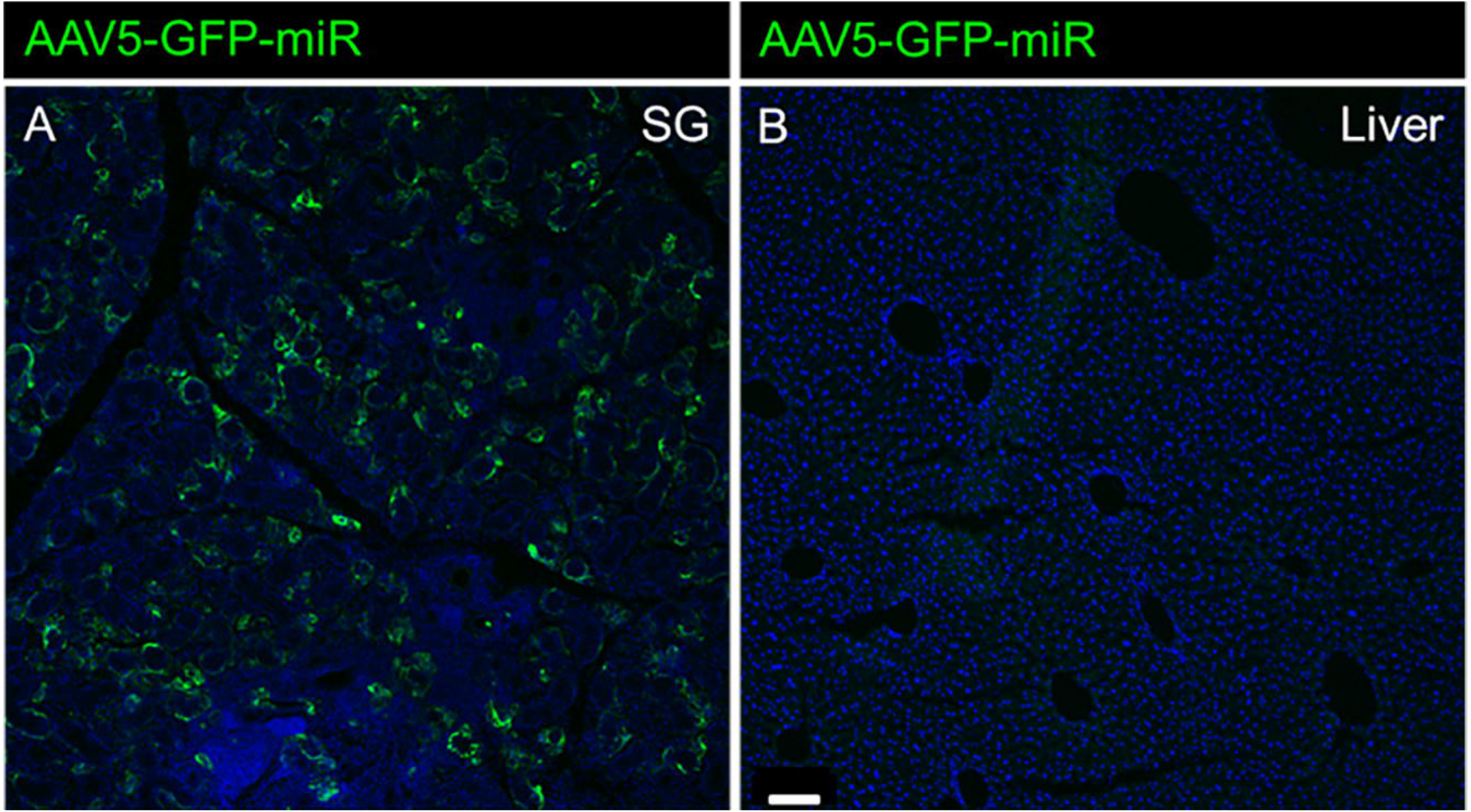
Negligible off target liver expression is observed with the inclusion of miR122 and miR206 TS into the AAV5 vector. GFP transduction in salivary glands (A) and liver (B) one month after vector administration was used as a marker of transduction.

APN KO mice with rescued salivary adiponectin expression were generated by administering 1×10^12^ vg of AAV5-APN-miR to the submandibular salivary glands of APN KO mice via ductile cannulation. AAV5-GFP-miR was injected into the salivary glands of APN KO mice as a negative control. A global adiponectin rescue model (positive control) was generated by administering 1×10^12^ vg of AAV8-APN systemically to APN KO mice via tail vein injection. One month after vector administration, we performed brief-access taste response testing for Intralipid, sucrose, and QHCL (Figure 8). We observed a modest, yet significant increase in the behavioral responses to Intralipid (p < 0.05), but not sucrose or QHCL, in the salivary adiponectin rescue mice compared to APN KO control mice (Figure 8A). The response of the global adiponectin rescue was not significantly different from that of the APN KO control mice for any of the tastants tested (Figure 8B).

**Figure 8.**
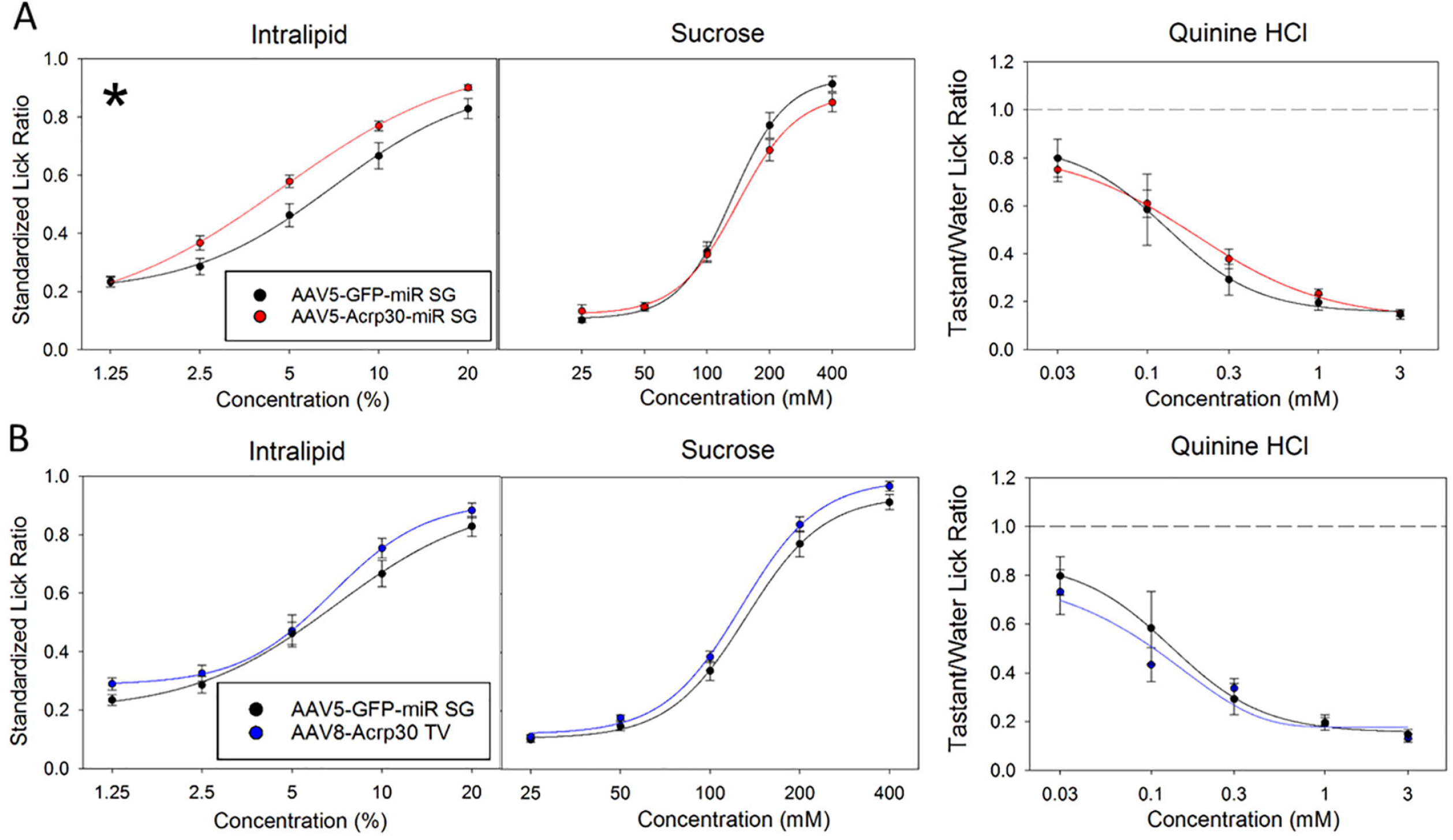
Behavioral taste response testing of salivary and global adiponectin rescue models. A) Brief-access taste response testing of salivary adiponectin rescue (red) and APN KO (black) for sucrose, Intralipid, and QHCL. B) Brief-access taste response testing of global adiponectin rescue (blue) and APN KO (black) for sucrose, Intralipid, and QHCL. *Significance in taste response was determined by two-way ANOVA and *post hoc* Holm-Sidak t-test with p < 0.05.

Finally, upon completion of behavioral testing, saliva and blood samples were drawn for adiponectin quantification by ELISA (Figure 9). As expected, plasma and saliva samples from APN KO mice were negative for adiponectin (Figure 9C). Mice receiving the AAV5-APN-miR vector show diminished circulating levels of adiponectin (11.66 ± 10.32 ng/ml; Figure 9A), approximately 1000-fold less than seen in WT mice (6.088 μg/ml; Figure 9C) and less likely to have a biological impact (Frühbeck *et al*. 2017). In saliva however, they express adiponectin at 3.94 ± 4.07 μg/ml (Figure 9A), similar to what is expected in WT mice (0.851 ng/ml; Figure 9C) and humans (Lin *et al*. 2014). By contrast, mice receiving systemic AAV8-APN showed much higher plasma levels of adiponectin (744.52 ± 365.67 μg/ml), but significantly lower levels in saliva (1.80 ± 0.83 ng/ml; Figure 9B).

**Figure 9.**
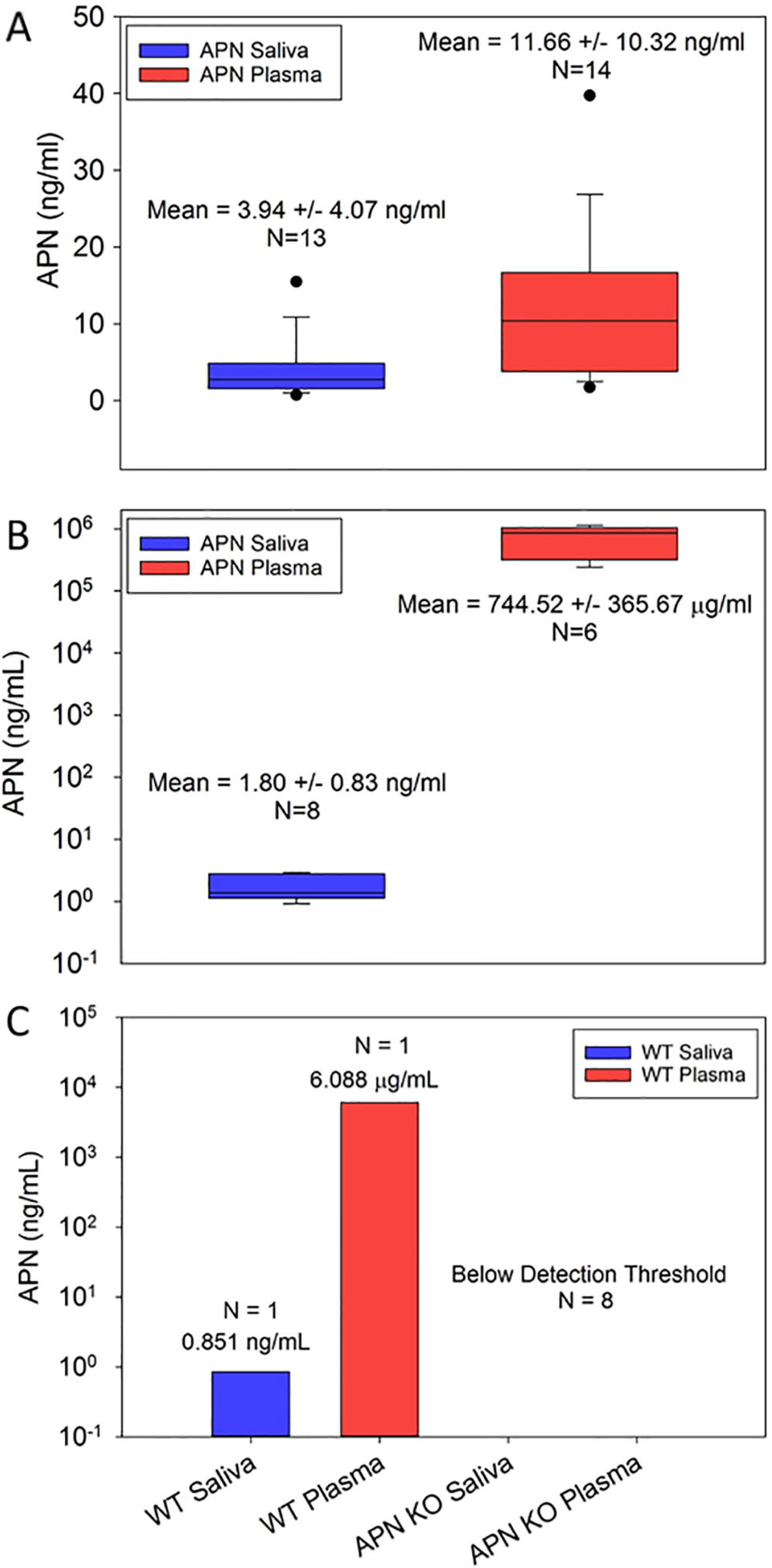
Quantification of adiponectin levels in rescue models by ELISA. A) Saliva (blue) and plasma (red) adiponectin levels of salivary adiponectin rescue mice. B) Saliva (blue) and plasma (red) adiponectin levels of global adiponectin rescue mice. C) Saliva (blue) and plasma (red) adiponectin levels in a single WT (C57BL/6) mouse, as well as confirmation of adiponectin loss in APN KO saliva and plasma.

## Discussion

TRCs and associated taste nerves express a diversity of receptors for peptide hormones related to the control of metabolism and satiety. The expression of two adiponectin receptors – Adipor1 and T-cadherin – in TRCs suggests an additional degree of complexity for modulation of TRC function by peptides acting as autocrine, paracrine, and/or endocrine factors. The expression patterns of different peptide receptors (as well as their peptide ligands) can vary significantly. For example, glucagon receptors are expressed in PLCβ2-positive Type 2 TRCs (Elson *et al*. 2010); oxytocin receptors are found in glial-like Type 1 cells (Sinclair *et al*. 2010); and the receptor for GLP-1 is localized to afferent nerve terminals innervating taste buds (Shin *et al*. 2008). We immunolocalized T-cadherin to a subset of PLCβ2-positive, G_α_-gustducin-positive TRCs, suggesting that adiponectin might affect responses to taste stimuli transduced by these signaling proteins. We also noted some T-cadherin-expressing cells that lacked immunostaining for PLCβ2, but also for markers of Type 1 and Type 3 TRCs. One possibility is that PLCβ2-negative, T-cadherin-positive TRCs represent an earlier stage of Type 2 cell differentiation and have not yet begun to express PLCβ2. Due to multiple antibody constraints (e.g., species compatibility), we were not able to fully resolve the exact expression patterns of each adiponectin receptor in taste buds using IHC alone. Transriptomic analyses of individual TRCs would be very useful for fully elucidating the expression profile of adiponectin receptors in TRCs.

While transcriptomic analysis of CV taste buds indicated that all three genes encoding canonical adiponectin receptors – *Adipor1, Adipor2*, and *Cdh13* – are expressed in TRCs, immunohistochemical staining showed that Adipor2 is excluded from TRCs and is instead localized to surrounding non-taste tissue. The discrepancy between the two techniques is not wholly surprising, as the taste buds used in the RNA-seq study were collected by manual dissection making low level contamination with non-taste tissue likely. Furthermore, differential localization of Adipor1 and Adipor2 in lingual tissue is consistent with observations in other tissues (Beylot *et al*. 2006).

Several peptides that affect blood glucose homeostasis, satiation, gastric emptying and secretion of digestive enzymes – including PYY, GLP-1 and glucagon (Batterham *et al*. 2002; Batterham *et al*. 2006; Hellström *et al*. 2004; Kieffer and Habener 1999; Nadkarni *et al*. 2014) – are produced in the oral cavity and impact taste responsiveness (Dotson *et al*. 2013; Elson *et al*. 2010; La Sala *et al*. 2013; Martin *et al*. 2009; Shin *et al*. 2008; Takai *et al*. 2015). While the majority of these peptides are produced in taste buds (Dotson *et al*. 2013), a few including leptin (Kawai *et al*. 2000), PYY (Acosta *et al*. 2011; La Sala *et al*. 2013), and oxytocin (Sinclair *et al*. 2010) are produced in distant tissues and likely reach the taste buds through saliva or the bloodstream. Adiponectin appears to fit this category, as well. This peptide has been widely studied because it plays critical roles in adipocyte metabolism, fatty acid oxidation, and insulin sensitivity (Lihn *et al*. 2005). Animal studies have shown that exogenous adiponectin administration leads to weight loss and insulin sensitization, and low levels of circulating adiponectin are correlated with metabolic syndrome in obese humans (Lin *et al*. 2007; Matafome *et al*. 2014; Shklyaev *et al*. 2003). However, adiponectin was not previously known to target the gustatory system.

Because the primary function of TRCs is to detect tastants and transduce this information to afferent gustatory nerve fibers, we reasoned that adiponectin may modulate taste responsiveness. Surprisingly, APN KO mice and their wildtype controls showed equivalent taste behavior responses to prototypical taste stimuli. However, the elimination of adiponectin may be insufficient to alter key downstream cellular signaling pathways in mice, as there may be compensatory mechanisms that are able to supplement for the lack of adiponectin (Ma *et al*. 2002). To address this issue, we performed the same behavioral testing in APN KO mice that had been rescued with rAAV-mediated expression of adiponectin either globally or specifically in salivary glands. Mice receiving the salivary adiponectin rescue (but not the global rescue mice or control APN KO mice) showed a modest but significant increase in behavioral responsiveness to Intralipid. Whether lipids elicit a distinct taste perceptual quality remains controversial, but they clearly can impact gustatory responses (Ozdener *et al*. 2014). Several putative “fat taste” receptors have been suggested in rodent taste buds, including the fatty acid translocase CD36 and the fatty acid-sensitive G protein-coupled receptor GPR120 (Cartoni *et al*. 2010). Interestingly, adiponectin has been reported to upregulate CD36 expression in cardiomyocytes via activation of the AMPK pathway (Chabowski *et al*. 2006; Fang *et al*. 2010). It is unclear whether a similar response may be present in TRCs.

Viral-mediated expression peptide hormones may be a useful strategy for modulating taste in a clinical context. By targeting expression to just the salivary glands, salivary adiponectin expression reached wildtype levels while circulating adiponectin levels were 1000-fold less than those in a WT mouse (Frühbeck *et al*. 2017). However, we were unable to completely limit adiponectin expression to either blood or saliva in either rescue model. For the salivary rescue model to limit off-target expression, we both directly injected AAV vectors driven by the chicken β-actin promoter into the salivary gland, and included micro RNAs for miR122 and miR206 which are liver and skeletal muscle specific, respectively. Even so, it was obvious that viral particles were still entering the circulation. Circulating adiponectin seen in this model could also be due to limited off-target transduction of non-salivary tissue, or the transduced salivary cells themselves may secrete adiponectin nonspecifically into both the blood and the saliva. In the global adiponectin rescue model, circulating adiponectin is likely transferred into saliva (Wang *et al*. 2013). Altogether, however, the salivary rescue model provided an impressive degree of expression control.

In summary we have shown that adiponectin receptors Adipor1 and T-cadherin are expressed in subpopulations of TRCs and that saliva-derived adiponectin can positively modulate taste behavioral responsiveness to Intralipid under certain conditions. A clearer understanding of the mechanisms by which adiponectin impacts TRC function awaits further studies of both oral lipid sensing and adiponectin-dependent signaling in the peripheral gustatory system.

## Acknowledgements

Thanks to the UF Center for Smell and Taste’s Chemosensory Behavior core for access to behavioral testing instruments. Thanks to Dr. Xiao-Rong Peng for their generous donation of both Adipor1 and Adipor2 antibodies.

## Financial Conflict of Interest

At the time of study, author C. D. Dotson was employed at the University of Florida. He has since been hired at the Coca Cola company, and has provided only editorial guidance since his hire at Coca Cola. This change of employment in no way influenced funding of the study or the results obtained. All other authors have no conflict of interests to declare.

## Funding Sources

NIH NIDCD F31 DC015751 and R01 DC012819

## References

Acosta A, Hurtado MD, Gorbatyuk O, La Sala M, Duncan D, Aslanidi G, Campbell-Thompson M, Zhang L, Herzog H, Voutetakis A, Baum BJ, Zolotukhin S. 2011. Salivary PYY: a putative bypass to satiety. PLoS One 6: e26137.

Awazawa M, Ueki K, Inabe K, Yamauchi T, Kubota N, Kaneko K, Kobayashi M, Iwane A, Sasako T, Okazaki Y, Ohsugi M, Takamoto I, Yamashita S, Asahara H, Akira S, Kasuga M, Kadowaki T. 2011. Adiponectin enhances insulin sensitivity by increasing hepatic IRS-2 expression via a macrophage-derived IL-6-dependent pathway. Cell Metab 13: 401–412.

Baquero AF, Gilbertson TA. 2011. Insulin activates epithelial sodium channel (ENaC) via phosphoinositide 3-kinase in mammalian taste receptor cells. Am J Physiol Cell Physiol 300: C860–871.

Batterham RL, Cowley MA, Small CJ, Herzog H, Cohen MA, Dakin CL, Wren AM, Brynes AE, Low MJ, Ghatei MA, Cone RD, Bloom SR. 2002. Gut hormone PYY(3–36) physiologically inhibits food intake. Nature 418: 650–654.

Batterham RL, Heffron H, Kapoor S, Chivers JE, Chandarana K, Herzog H, Le Roux CW, Thomas EL, Bell JD, Withers DJ. 2006. Critical role for peptide YY in protein-mediated satiation and body-weight regulation. Cell Metab 4: 223–233.

Beylot M, Pinteur C, Peroni O. 2006. Expression of the adiponectin receptors AdipoR1 and AdipoR2 in lean rats and in obese Zucker rats. Metabolism 55: 396–401.

Bjursell M, Ahnmark A, Bohlooly-Y M, William-Olsson L, Rhedin M, Peng XR, Ploj K, Gerdin AK, Arnerup G, Elmgren A, Berg AL, Oscarsson J, Lindén D. 2007. Opposing effects of adiponectin receptors 1 and 2 on energy metabolism. Diabetes 56: 583–593.

Bobbert T, Rochlitz H, Wegewitz U, Akpulat S, Mai K, Weickert MO, Möhlig M, Pfeiffer AF, Spranger J. 2005. Changes of adiponectin oligomer composition by moderate weight reduction. Diabetes 54: 2712–2719.

Cartoni C, Yasumatsu K, Ohkuri T, Shigemura N, Yoshida R, Godinot N, le Coutre J, Ninomiya Y, Damak S. 2010. Taste preference for fatty acids is mediated by GPR40 and GPR120. J Neurosci 30: 8376–8382.

Chabowski A, Momken I, Coort SL, Calles-Escandon J, Tandon NN, Glatz JF, Luiken JJ, Bonen A. 2006. Prolonged AMPK activation increases the expression of fatty acid transporters in cardiac myocytes and perfused hearts. Mol Cell Biochem 288: 201–212.

Chaudhari N, Roper SD. 2010. The cell biology of taste. J Cell Biol 190: 285–296.

Costes SV, Daelemans D, Cho EH, Dobbin Z, Pavlakis G, Lockett S. 2004. Automatic and quantitative measurement of protein-protein colocalization in live cells. Biophys J 86: 3993–4003.

Denzel MS, Scimia MC, Zumstein PM, Walsh K, Ruiz-Lozano P, Ranscht B. 2010. T-cadherin is critical for adiponectin-mediated cardioprotection in mice. J Clin Invest 120: 4342–4352.

Ding C, Li L, Su YC, Xiang RL, Cong X, Yu HK, Li SL, Wu LL, Yu GY. 2013. Adiponectin increases secretion of rat submandibular gland via adiponectin receptors-mediated AMPK signaling. PLoS One 8: e63878.

Dotson CD, Geraedts MC, Munger SD. 2013. Peptide regulators of peripheral taste function. Semin Cell Dev Biol 24: 232–239.

Elson AE, Dotson CD, Egan JM, Munger SD. 2010. Glucagon signaling modulates sweet taste responsiveness. FASEB J 24: 3960–3969.

Fang X, Palanivel R, Cresser J, Schram K, Ganguly R, Thong FS, Tuinei J, Xu A, Abel ED, Sweeney G. 2010. An APPL1-AMPK signaling axis mediates beneficial metabolic effects of adiponectin in the heart. Am J Physiol Endocrinol Metab 299: E721–729.

Frühbeck G, Catalán V, Rodríguez A, Ramírez B, Becerril S, Portincasa P, Gómez-Ambrosi J. 2017. Normalization of adiponectin concentrations by leptin replacement in ob/ob mice is accompanied by reductions in systemic oxidative stress and inflammation. Sci Rep 7: 2752.

Geisler A, Schön C, Größl T, Pinkert S, Stein EA, Kurreck J, Vetter R, Fechner H. 2013. Application of mutated miR-206 target sites enables skeletal muscle-specific silencing of transgene expression of cardiotropic AAV9 vectors. Mol Ther 21: 924–933.

Glendinning JI, Gresack J, Spector AC. 2002. A high-throughput screening procedure for identifying mice with aberrant taste and oromotor function. Chem Senses 27: 461–474.

Grimm D, Kern A, Rittner K, Kleinschmidt JA. 1998. Novel tools for production and purification of recombinant adenoassociated virus vectors. Hum Gene Ther 9: 2745–2760.

Hellström PM, Geliebter A, Näslund E, Schmidt PT, Yahav EK, Hashim SA, Yeomans MR. 2004. Peripheral and central signals in the control of eating in normal, obese and binge-eating human subjects. Br J Nutr 92 Suppl 1: S47–57.

Huang YA, Maruyama Y, Stimac R, Roper SD. 2008. Presynaptic (Type III) cells in mouse taste buds sense sour (acid) taste. J Physiol 586: 2903–2912.

Hurtado MD, Acosta A, Riveros PP, Baum BJ, Ukhanov K, Brown AR, Dotson CD, Herzog H, Zolotukhin S. 2012. Distribution of Y-receptors in murine lingual epithelia. PLoS One 7: e46358.

Katano H, Kok MR, Cotrim AP, Yamano S, Schmidt M, Afione S, Baum BJ, Chiorini JA. 2006. Enhanced transduction of mouse salivary glands with AAV5-based vectors. Gene Ther 13: 594–601.

Katsiougiannis S, Kapsogeorgou EK, Manoussakis MN, Skopouli FN. 2006. Salivary gland epithelial cells: a new source of the immunoregulatory hormone adiponectin. Arthritis Rheum 54: 2295–2299.

Katsiougiannis S, Tenta R, Skopouli FN. 2010. Activation of AMP-activated protein kinase by adiponectin rescues salivary gland epithelial cells from spontaneous and interferon-gamma-induced apoptosis. Arthritis Rheum 62: 414–419.

Kawai K, Sugimoto K, Nakashima K, Miura H, Ninomiya Y. 2000. Leptin as a modulator of sweet taste sensitivities in mice. Proc Natl Acad Sci U S A 97: 11044–11049.

Kieffer TJ, Habener JF. 1999. The glucagon-like peptides. Endocr Rev 20: 876–913.

La Sala MS, Hurtado MD, Brown AR, Bohórquez DV, Liddle RA, Herzog H, Zolotukhin S, Dotson CD. 2013. Modulation of taste responsiveness by the satiation hormone peptide YY. FASEB J 27: 5022–5033.

Lihn AS, Pedersen SB, Richelsen B. 2005. Adiponectin: action, regulation and association to insulin sensitivity. Obes Rev 6: 13–21.

Lin H, Maeda K, Fukuhara A, Shimomura I, Ito T. 2014. Molecular expression of adiponectin in human saliva. Biochem Biophys Res Commun 445: 294–298.

Lin HV, Kim JY, Pocai A, Rossetti L, Shapiro L, Scherer PE, Accili D. 2007. Adiponectin resistance exacerbates insulin resistance in insulin receptor transgenic/knockout mice. Diabetes 56: 1969–1976.

Ma K, Cabrero A, Saha PK, Kojima H, Li L, Chang BH, Paul A, Chan L. 2002. Increased beta - oxidation but no insulin resistance or glucose intolerance in mice lacking adiponectin. J Biol Chem 277: 34658–34661.

Martin B, Dotson CD, Shin YK, Ji S, Drucker DJ, Maudsley S, Munger SD. 2009. Modulation of taste sensitivity by GLP-1 signaling in taste buds. Ann N Y Acad Sci 1170: 98–101.

Matafome P, Rodrigues T, Pereira A, Letra L, Azevedo H, Paixão A, Silvério M, Almeida A, Sena C, Seiça R. 2014. Long-term globular adiponectin administration improves adipose tissue dysmetabolism in high-fat diet-fed Wistar rats. Arch Physiol Biochem 120: 147–157.

Mbiene JP, Roberts JD. 2003. Distribution of keratin 8-containing cell clusters in mouse embryonic tongue: evidence for a prepattern for taste bud development. J Comp Neurol 457: 111–122.

Ming D, Ninomiya Y, Margolskee RF. 1999. Blocking taste receptor activation of gustducin inhibits gustatory responses to bitter compounds. Proc Natl Acad Sci U S A 96: 9903–9908.

Miyoshi MA, Abe K, Emori Y. 2001. IP(3) receptor type 3 and PLCbeta2 are co-expressed with taste receptors T1R and T2R in rat taste bud cells. Chem Senses 26: 259–265.

Nadkarni P, Chepurny OG, Holz GG. 2014. Regulation of glucose homeostasis by GLP-1. Prog Mol Biol Transl Sci 121: 23–65.

Ozdener MH, Subramaniam S, Sundaresan S, Sery O, Hashimoto T, Asakawa Y, Besnard P, Abumrad NA, Khan NA. 2014. CD36- and GPR120-mediated Ca^2^□ signaling in human taste bud cells mediates differential responses to fatty acids and is altered in obese mice. Gastroenterology 146: 995–1005.

Piedra J, Ontiveros M, Miravet S, Penalva C, Monfar M, Chillon M. 2015. Development of a rapid, robust, and universal picogreen-based method to titer adeno-associated vectors. Hum Gene Ther Methods 26: 35–42.

Shigemura N, Iwata S, Yasumatsu K, Ohkuri T, Horio N, Sanematsu K, Yoshida R, Margolskee RF, Ninomiya Y. 2013. Angiotensin II modulates salty and sweet taste sensitivities. J Neurosci 33: 6267–6277.

Shin YK, Martin B, Golden E, Dotson CD, Maudsley S, Kim W, Jang HJ, Mattson MP, Drucker DJ, Egan JM, Munger SD. 2008. Modulation of taste sensitivity by GLP-1 signaling. J Neurochem 106: 455–463.

Shklyaev S, Aslanidi G, Tennant M, Prima V, Kohlbrenner E, Kroutov V, Campbell-Thompson M, Crawford J, Shek EW, Scarpace PJ, Zolotukhin S. 2003. Sustained peripheral expression of transgene adiponectin offsets the development of diet-induced obesity in rats. Proc Natl Acad Sci U S A 100: 14217–14222.

Sinclair MS, Perea-Martinez I, Dvoryanchikov G, Yoshida M, Nishimori K, Roper SD, Chaudhari N. 2010. Oxytocin signaling in mouse taste buds. PLoS One 5: e11980.

Takai S, Yasumatsu K, Inoue M, Iwata S, Yoshida R, Shigemura N, Yanagawa Y, Drucker DJ, Margolskee RF, Ninomiya Y. 2015. Glucagon-like peptide-1 is specifically involved in sweet taste transmission. FASEB J 29: 2268–2280.

Vandenbeuch A, Anderson CB, Parnes J, Enjyoji K, Robson SC, Finger TE, Kinnamon SC. 2013. Role of the ectonucleotidase NTPDase2 in taste bud function. Proc Natl Acad Sci U S A 110: 14789–14794.

Villarreal-Molina MT, Antuna-Puente B. 2012. Adiponectin: anti-inflammatory and cardioprotective effects. Biochimie 94: 2143–2149.

Wang J, Liang Y, Wang Y, Cui J, Liu M, Du W, Xu Y. 2013. Computational prediction of human salivary proteins from blood circulation and application to diagnostic biomarker identification. PLoS One 8: e80211.

Yamauchi T, Kamon J, Minokoshi Y, Ito Y, Waki H, Uchida S, Yamashita S, Noda M, Kita S, Ueki K, Eto K, Akanuma Y, Froguel P, Foufelle F, Ferre P, Carling D, Kimura S, Nagai R, Kahn BB, Kadowaki T. 2002. Adiponectin stimulates glucose utilization and fatty-acid oxidation by activating AMP-activated protein kinase. Nat Med 8: 1288–1295.

Yee CL, Yang R, Böttger B, Finger TE, Kinnamon JC. 2001. “Type III” cells of rat taste buds: immunohistochemical and ultrastructural studies of neuron-specific enolase, protein gene product 9.5, and serotonin. J Comp Neurol 440: 97–108.

Yoon MJ, Lee GY, Chung JJ, Ahn YH, Hong SH, Kim JB. 2006. Adiponectin increases fatty acid oxidation in skeletal muscle cells by sequential activation of AMP-activated protein kinase, p38 mitogen-activated protein kinase, and peroxisome proliferator-activated receptor alpha. Diabetes 55: 2562–2570.

Zhao FL, Shen T, Kaya N, Lu SG, Cao Y, Herness S. 2005. Expression, physiological action, and coexpression patterns of neuropeptide Y in rat taste-bud cells. Proc Natl Acad Sci U S A 102: 11100–11105.

Zolotukhin S. 2013. Metabolic hormones in saliva: origins and functions. Oral Dis 19: 219–229.

Zolotukhin S, Byrne BJ, Mason E, Zolotukhin I, Potter M, Chesnut K, Summerford C, Samulski RJ, Muzyczka N. 1999. Recombinant adeno-associated virus purification using novel methods improves infectious titer and yield. Gene Ther 6: 973–985.

Zolotukhin S, Potter M, Hauswirth WW, Guy J, Muzyczka N. 1996. A “humanized” green fluorescent protein cDNA adapted for high-level expression in mammalian cells. J Virol 70: 4646–4654.

Zolotukhin S, Potter M, Zolotukhin I, Sakai Y, Loiler S, Fraites TJ, Chiodo VA, Phillipsberg T, Muzyczka N, Hauswirth WW, Flotte TR, Byrne BJ, Snyder RO. 2002. Production and purification of serotype 1, 2, and 5 recombinant adeno-associated viral vectors. Methods 28: 158–167.

